# Odour representations supporting ethology-relevant categorisation and discrimination in the *Drosophila* mushroom body

**DOI:** 10.1101/2025.01.25.634657

**Authors:** Ivy Chi Wai Chan, Felipe Yaroslav Kalle Kossio, Gaia Tavosanis

## Abstract

Neural representations of sensory stimuli serve multiple distinct purposes, from the rapid recognition of familiar environments, to the precise identification of individual salient cues. In the insect mushroom body (MB), odours are encoded by the activity of Kenyon cells (KCs). The random wiring of olfactory projection neurons (PNs) and KCs in the MB calyx is thought to enhance odour discrimination. Here, we examined the impact of deviations from random wiring and demonstrated their significant roles in shaping odour representations. We confirm that different KC types have distinct PN input biases correlated with the contextual relevance of the odour information delivered by the PNs. By recording the functional responses of different KC types to ethologically defined odour categories, we found that the αβ and α’β’ KCs produce segregated representations of relevant odour groups, potentially enhancing the categorisation of odours based on ethological relevance. Simultaneously, these same KC types displayed distinct representations for food-related odours, supporting precise discrimination. In contrast, γ KCs lacked significant segregation of odour representations by ethological category. Computational simulations refined with our functional data indicated that the specific PN input connection pattern of individual KC types largely accounts for the observed representations. Taken together, we propose that individual KC types process odour information with distinct objectives, supporting both ethological categorisation and discrimination.

## Introduction

In their natural environment, animals are continuously exposed to complex sensory information. Survival often depends on their ability to rapidly filter through this information, recognizing relevant signal categories to evade dangers and locate essential resources like food or mates. Conversely, many species demonstrate remarkable precision in identifying and assigning value to specific sensory stimuli, effectively differentiating them from similar ones. These apparently contrasting ways of evaluating sensory information are essential for instance to allow foraging insects to recognize individual flowers and avoid re-visiting them, while generalising across similar types to maximise efficiency ^1–3^. Thus, the ability to perform both stimulus discrimination and categorisation is vital for interacting efficiently with a complex sensory environment. However, the mechanisms by which the brain’s higher processing centres reconcile the potentially conflicting computations performed on common sensory inputs—distinguishing unique versus shared traits—remain a compelling puzzle in neuroscience.

Stimulus discrimination is often computed within specialised brain circuits known to facilitate distinct representations of similar inputs ^4,5^, such as the mammalian cerebellum and dentate gyrus, and the *Drosophila* mushroom body (MB), which is the centre for associative learning ^6,7^. These structures exhibit a specific organisational motif whereby a relatively small number of neurons in an input layer connect to a much larger number of neurons in an expansion layer; together with feedforward or feedback inhibition, this organization supports sparse, decorrelated and high dimensional responses that enhance pattern separation ^8,9,4^. On the contrary, how dissimilar odours can be recognized as a category (e.g. food) remains less well-understood. In zebrafish, for instance, the temporal pattern of mitral cells’ spikes - resembling olfactory projection neurons in insects - carry distinct information about both odour identity and category, which can be differentially interpreted by various target neurons^10^.

With a rich set of genetic tools and two fully-annotated connectomes of the olfactory pathway ^11–13^, the *Drosophila* MB serves as an important model to study how circuit architecture can define the functional output of a pattern separation circuit ^14,15^. In *Drosophila*, olfactory information acquired by olfactory sensory neurons (OSNs) is delivered to the antennal lobe (AL), which is segregated into 50 glomeruli based on the receptor types expressed by the OSNs. Uniglomerular second order projection neurons (PNs) then transfer information from their respective AL glomerulus to the MB input region, the calyx, where they form large axonal boutons. In the calyx, around 2000 Kenyon cells (KCs) receive olfactory inputs from the 50 types of PNs, yielding a 40-fold circuit expansion (Fig. 1a). Individual PN boutons are enveloped by the dendritic claws of on average 12 KCs, forming large synaptic complexes called microglomeruli ^16–18^. Each KC connects via its few dendritic claws to distinct PN boutons ^14,19,20^, thus receiving a mix of inputs in the presence of inhibitory signals from the anterior paired lateral (APL) neuron ^21,22^. The PN-to-KC connection pattern in the MB calyx plays a key role in the formation of sparse, high-dimensional KC responses to odour stimulation ^19,15,23^. Three main types of KCs, αβ, α’β’ and γ display distinct roles in the formation, processing and retrieval of memory ^24–29^, and most PN boutons connect to all three KC types within microglomeruli ^17^.

**Figure 1.**
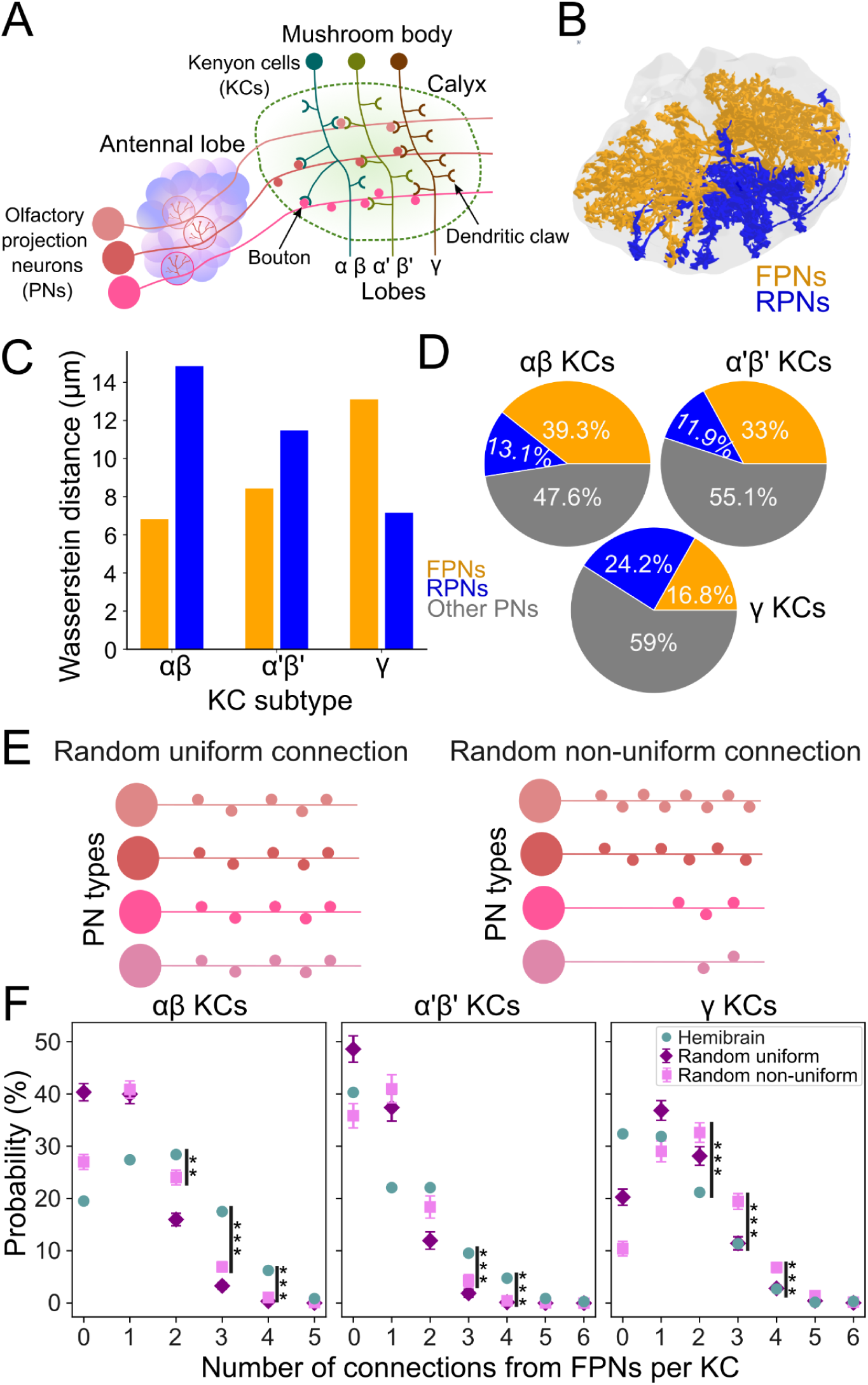
Structured spatial organisation of PN to KC connections in the calyx reflects ethological odour categories. (a) In the MB calyx, over 2000 KCs receive olfactory inputs from 50 types of uniglomerular olfactory PNs. KCs are subdivided into three main types based on their origin and their axonal projections into the α and β, α’ and β’ or γ lobes. Each KC has multiple dendritic claws: on average 5 for αβ, 4 for α’β’, and 8 for γ KCs. Each claw receives input from an individual PN bouton. Hence, each KC receives a combination of inputs from multiple PNs. (b) Reconstructed FPNs’ (orange) and RPNs’ (blue) axonal collaterals from the Hemibrain dataset in the calycal volume (grey). The FPNs distribute their collaterals in the ventral-posterior region of the calyx, while the RPNs in the dorsal-anterior region. (c) Wasserstein distance between presynapse distribution of FPNs/RPNs (orange/blue) and postsynapse distributions of each KC type. (d) Fractions of synaptic inputs received from FPNs (orange), RPNs (blue) and other PNs (grey) by αβ KCs, α’β’ KCs or γ KCs. αβ and α’β’ KCs sample the FPNs more extensively than γ KCs. Conversely, γ KCs sample RPNs more extensively than αβ and α’β’ KCs. (e) In the *random-uniform* connection model, each KC claw has an equal chance of connecting to one of the 50 PN types. In the *random non-uniform* connection model, each KC claw connects to a PN type with probability equal to the fraction of connections to that PN type in the Hemibrain dataset. In both models, the number of claws of each KC was drawn from the claw number distribution for the corresponding KC type derived from the Hemibrain dataset. (f) KCs to FPNs connection probabilities in Hemibrain or in the two random connections models. The graphs show the probability that a KC forms a certain number of connections with FPNs in the Hemibrain (turquoise), in the random uniform (purple) and random non-uniform (pink) connection models. * p<0.05, ** p<0.01, *** p<0.001, Kruskal-Wallis H-test.

Previous theoretical work, supported by experimental evidence, had indicated that close to random PN bouton to KC claw connections result in high-dimensional odour representations and good discrimination performance ^14,30^. Nonetheless, recent electron-microscopy based connectome data showed that some PN types connect in a deviating-from-random fashion to KCs ^12,20^. In particular, in the Female Adult Fly Brain dataset (FAFB), PNs that deliver information about food-related odours are more likely to be connected to the same individual αβ or α’β’ KCs than expected from a random distribution ^20^. This raised the hypothesis that the MB calyx circuit, in addition to performing odour discrimination, supports odour categorisation based on ethologically relevant criteria. Furthermore, it questions whether different KC types might differently process odour information delivered by the PNs, for example favoring discrimination or categorisation of some odour categories. However, functional evidence addressing these points is lacking.

Here, we combined connectome analysis, functional imaging and network modelling to investigate how the anatomy of the *Drosophila* MB calyx supports extracting specific information aspects from odour stimuli. We present evidence that the MB, in addition to odour discrimination, performs odour categorisation of ethologically relevant odour groups. Our data indicate that the three KC types perform distinct operations as they sample the olfactory space differently, deviating from the hypothesised random sampling, and yet supporting efficient discrimination.

## Results

### Distinct sampling biases of the three KC types based on the ethological category of odours

To elucidate how odours of different ethological relevance are represented in the MB, we analysed the projection patterns and connection logic between PNs and KC types in the calyx, utilising data from the Hemibrain connectome dataset ^12,31^. Most AL glomeruli show strong responses to odours with specific behavioural relevance such as feeding, reproduction, and danger ^32^. Among those that respond to various food-related odours, their corresponding PN types converge more often than by chance to αβ and α’β’ KCs in the FAFB dataset ^20^. These food-odour-responding PNs (FPNs, see Methods) are specialised in detecting yeast-related odours and by-products of their alcoholic fermentation ^32^. In this study, we focused our analysis on these FPNs and compared them to PNs that respond to various pheromones and odours present at egg-laying sites, termed reproduction-odour-responding PNs (RPNs, see Methods).

In the Hemibrain dataset, the boutons from FPNs and RPNs were spatially segregated within the calyx (Fig. 1b). We found that the αβ and α’β’ KCs’ postsynapse distributions were spatially closer to the FPNs’ presynapse distribution than to the RPNs’ (Wasserstein distance, see Methods), while the γ KCs’ postsynapse distribution was closer to the RPNs’ presynapse distribution than to the FPNs’ (Fig. 1c). In agreement with this, the fraction of synapses that αβ and α’β’ KCs received from FPNs was larger (39.3 % and 33 %, respectively) compared to γ KCs (16.8 %, Fig. 1d). The distribution of PN-to-KC connections, that we define as synaptic contacts between a PN bouton and a KC claw, irrespective of the number of synapses formed, followed a similar pattern (Supplementary Fig. 1e). In contrast, the fraction of synapses that γ KCs received from RPNs was larger (24.2 %) compared to αβ and α’β’ KCs (13.1 % and 11.9 %, respectively, Fig. 1d).

To clarify the impact of the observed connection bias of αβ and α’β’ KCs towards FPNs, we generated two computational models in which the PN-to-KC connections were randomly formed and compared those to the Hemibrain dataset. For both models, the distribution of the number of claws formed by individual KC was inferred from the Hemibrain dataset for each KC type. In the *random uniform* connection model, each KC claw had an equal chance of connecting to one of the 50 PN types (Fig. 1e). Because the numbers of boutons formed in the calyx by different PN types are different ^20^, with FPNs displaying in general a higher number of boutons, we also generated a r*andom non-uniform* connection model. Here, each KC claw is connected to a PN type with probability proportional to the actual fraction of connections formed by that PN type in the Hemibrain dataset (Supplementary Table 1). This takes into account the different number of boutons per PN type and other possible factors.

Comparing the different connection logics, we found that αβ and α’β’ KCs exhibited a higher probability of forming multiple connections with FPNs than predicted by either random model (Fig. 1f). For instance, the probability of an αβ KC forming exactly three connections with FPNs was significantly higher in the Hemibrain dataset (17.5 %) compared to the random non-uniform (7.0 %) and random uniform (3.2 %) models. α’β’ KCs also showed a higher probability of forming three connections with FPNs (9.6 %) in the Hemibrain dataset compared to both random models (4.1 % and 1.9 %, respectively). For γ KCs, the number of connections with FPNs in the Hemibrain dataset deviated significantly from the random non-uniform connection model and resembled the random uniform connection model more closely. This suggested a negative bias of γ KCs towards the FPNs. Conversely, the number of connections with RPNs formed by individual KCs showed a negative bias for αβ and α’β’ KCs and a positive bias for γ KCs (Supplementary Fig. 1d).

Taken together, αβ and α’β’ KCs display a connection bias towards FPNs while γ KCs display a connection bias towards RPNs in the Hemibrain dataset, confirming and extending findings from the FAFB dataset ^20^. Furthermore, the connection bias of FPNs towards αβ and α’β’ KCs cannot be simply explained by random sampling that takes into account the different numbers of connections per PN type.

### Ethological odour category representation by KCs

To investigate the functional significance of the PN-to-KC type connection biases, we conducted *in-vivo* calcium volumetric imaging of all KC types simultaneously, as well as separately for each KC type, recording their somatic responses to a panel of 12 monomolecular odours from three ethological groups: food-, reproduction- and danger-related. The selected food-related odours consisted of esters derived from fruits and yeast fermentation, while reproduction-related odours included molecularly distinct pheromones as well as odours present at egg-laying sites. Danger-related odours, on the other hand, are aversive, signalling potential threats (Table 1). Each trial consisted of a series of 10 to 12 odours presented in a pseudo-random order for 2 seconds each with a 45-second interval between presentations (Fig. 2a). For each fly, we carried out 2 to 4 trials, allowing a 10-minute rest interval between them.

**Figure 2.**
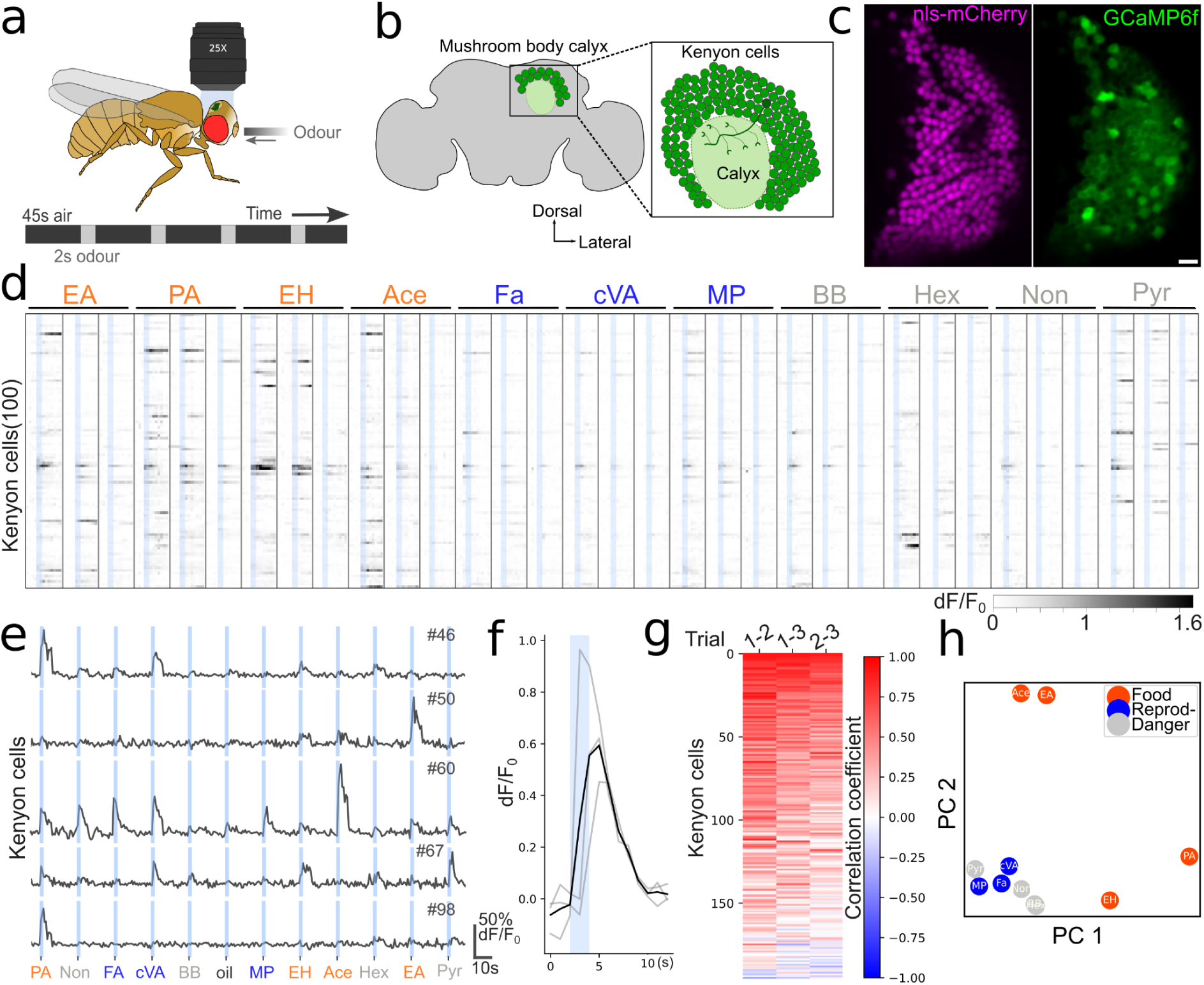
Distinct functional representations of odours of different categories in KCs. (a) A female fly expressing GCaMP6f and NLS-mCherry in all KCs or in a certain KC type was immobilised to an imaging chamber, with cuticle and fat bodies removed on the posterior side of the head, exposing the KCs. The fly was then placed under a 25X immersion objective at a two-photon microscope. Odour delivery was conducted via a nozzle positioned in front of the fly’s antennae, with each odour presented for 2 seconds (grey) followed by a 45-second interval of clean air flow (black). The sequence of odours was administered in a pseudo-random order. (b) An imaging plane (inset) contained 150-200 KCs cell bodies surrounding the calyx. (c) An example imaging plane averaged across time frames from an all-KCs fly (all data presented in c - g are from the same fly). The NLS-mCherry (magenta) localised in the nuclei of the KCs provided the structural labelling for motion correction while the GCaMP6f (green) in the cytoplasm allowed measurement of the calcium activity. (Scale bar = 10µm) (d) Normalised changes in fluorescence levels for each KC (rows) across time in response to odours. The blue shaded region indicates the 2 seconds odour delivery period. Each column corresponds to one trial and the three trials per odour are grouped. Food-related odours are in orange, reproduction-related odours in blue, and danger-related odours in red, see Table 1. (e) Normalised changes in fluorescence levels from five individual cells for a panel of 12 odours in a single trial. The blue shaded region indicates the odour delivery period. (f) Normalised changes in fluorescence levels (grey lines) of a single KC across three exposures to the odour Farnesol and averaged trace (black line). The blue shaded region indicates the odour delivery period. (g) Inter-trial correlation of odour-evoked responses for individual KCs (rows). Each column shows the correlation between different trials. (h) Principal component analysis of KC responses to a panel of 12 odours for all KCs (n = 3 flies, no. of KCs = 504).

**Table 1.** The panel of 12 monomolecular odours.

| **Ethological relevance** | **Odours** | **Abbreviation** | **Dilution** | **Primarily Activated Glomeruli** | **Reference** |
| --- | --- | --- | --- | --- | --- |
| Food | Pentyl acetate | PA | 1:100 | DM3, VC3 | <sup>32</sup> |
| Food | Ethyl hexanoate | EH | 1:100 | DM3, DM6 | <sup>32</sup> |
| Food | Ethyl acetate | EA | 1:100 | DM1, VM7 | <sup>32</sup> |
| Food | Acetone | Ace | 1:100 | DM1, DM4 | <sup>32</sup> |
| Egg-laying site / Food | Farnesol | Fa | 1:100 | DC3 | <sup>33</sup> |
| Pheromone | Methyl palmitate | MP | 1:100 | VA1d | <sup>34</sup> |
| Pheromone | Methyl laurate | ML | 1:100 | VA1v | <sup>34</sup> |
| Pheromone | 11-*cis*-vaccenyl acetate | cVA | 1:100 | DA1, DL3 | <sup>35–37</sup> |
| Danger | Butyl butyrate | BB | 1:100 | DC1, DM2 | <sup>38</sup> |
| Danger | 1-hexanol | Hex | 1:100 | VA3, VC3 | <sup>32,38</sup> |
| Danger | 2-nonanone | Non | 1:100 | D, VC3 | <sup>32</sup> |
| Danger | Pyrrolidine | Pyr | 1:1000 | VL1, VL2a | <sup>32</sup> |
| Control | Mineral oil | Oil |  |  |  |

We expressed cytoplasmic GCaMP6f to monitor odour-evoked calcium activity, together with nuclear NLS-mCherry, which provided structural labelling for motion correction and cell segmentation in *OK107-Gal4* (all-KCs, n = 3) or in specific KC types: *NP3061-Gal4* (αβ KCs, n = 3), *c305a-Gal4* (α’β’ KCs, n = 3), and *GMR71G10-Gal4* (γ KCs, n = 3) (Fig. 2b, c).

Throughout the sessions, we reliably tracked 504 cells from the all-KCs flies, and 290, 269 and 283 cells from flies expressing *GCaMP6f* and *NLS-mCherry* in αβ, α’β’ or γ KCs, respectively. Examples of KC responses to the odour panel from an all-KCs fly are shown in Fig. 2d and 2e. Consistent odour-evoked responses were observed across multiple sessions (Fig. 2f and 2g). The odour-evoked average activity for each KC was calculated by averaging across all sessions and across time (from odour onset to 5 seconds post-onset). The analysis included all KCs, regardless of their level of activity. All KCs responses from flies of the same genotype were pooled together (individual fly data are shown in Supplementary Figs. 3 - 6).

To explore the organisation of olfactory representations, we used this odour-evoked average activity for each KC in the all-KCs experiments to construct an odour representation vector for each odour, and projected these vectors to a lower-dimensional space using principal component analysis (PCA, Fig. 2h). Strikingly, we found that the representation of food-related odours clearly segregated from that of reproduction- or danger-related odours, already for the first two PCs. In addition, the Euclidean distances among representations of individual food-related odours were higher than among reproduction- and danger-related odours (Supplementary Fig. 3), despite a higher chemical similarity among the food-related odours than among the danger-related odours (Supplementary Fig. 2). Thus, KCs representations can support categorisation of food-related odours from other odours, and at the same time potentially preserve the capacity to discriminate among them.

### Segregation of food-related olfactory representations in αβ and α’β’ KCs

Given the biased connectivity of αβ and α’β’ KCs towards FPNs, we hypothesise that the higher capacity of KCs to segregate food-related odours from others and discriminate between them might be primarily contributed by αβ and α’β’ KCs. We therefore analysed the odour-evoked activity of individual KC types (αβ, α’β’, and γ), and used the average responses of individual cells to generate odour representation vectors that we projected to a lower-dimensional space using PCA (Fig. 3a). αβ and α’β’ KCs representations of food-related odours were segregated from the reproduction-related odour representations (Fig. 3a). In contrast, for γ KC, no clear segregation was observed between the two representations (Fig. 3a). This data suggested possible grouping of odour representations based on ethological significance within the activity space of αβ and α’β’ KCs, but not of γ KCs.

**Figure 3.**
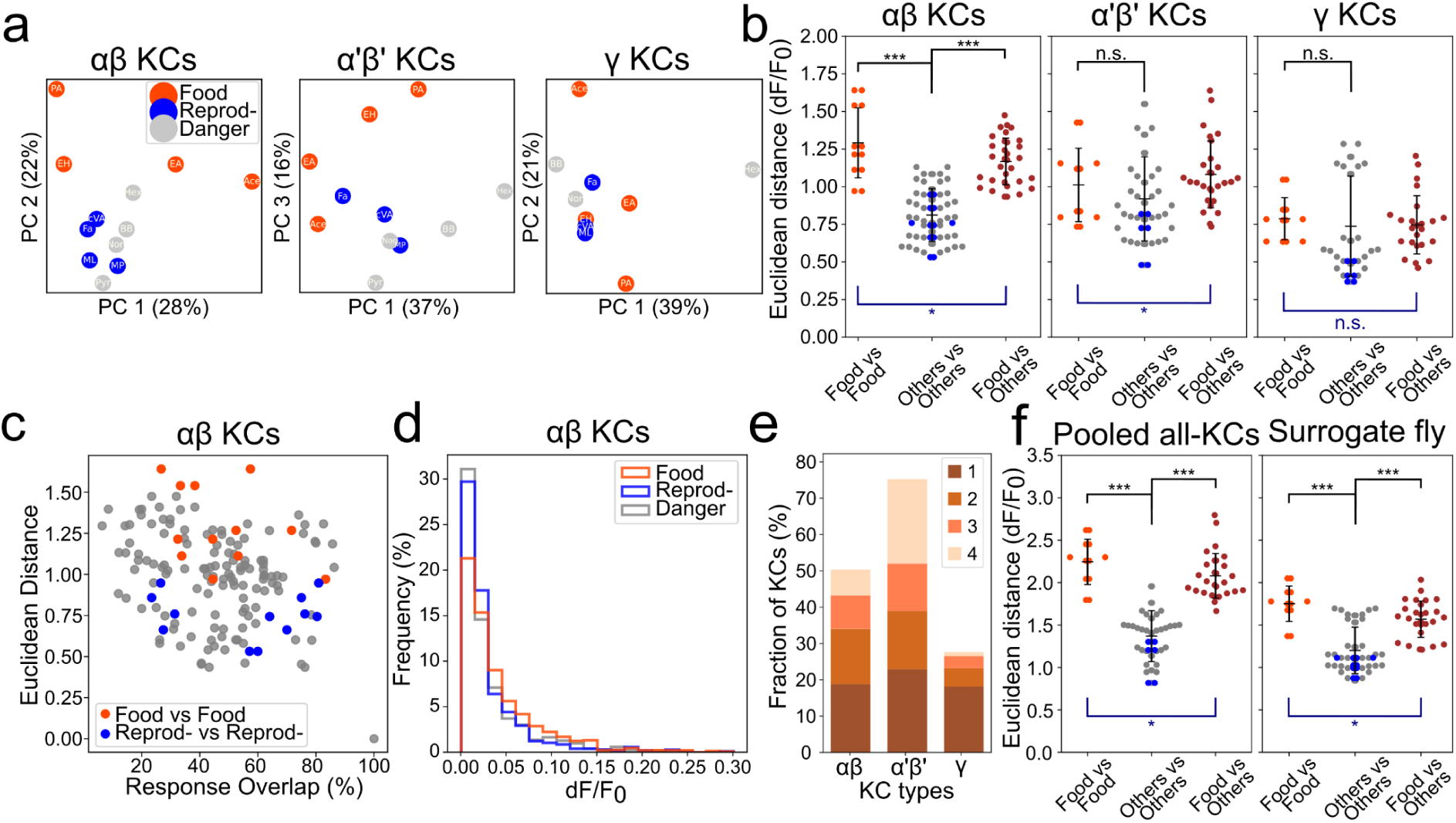
Ethology-based segregation of olfactory representation in αβ and α’β’ KCs. (a) PCA of KCs’ responses to a panel of 12 odours for each KC type: pooled αβ KCs flies (n = 3, 290 KCs), pooled α’β’ KCs flies (n = 3, 269 KCs) and pooled γ KCs flies (n = 3, 283 KCs). Food-related odours are shown in orange, reproduction-related odours in blue, and danger-related odours in red. The representations of food-related odours for αβ KCs and for α’β’ KCs were segregated from the rest. Conversely, in γ KCs, the odour representations of different groups are less segregated. (b) Euclidean distances between odour representations for the three KC types: pooled αβ KCs, pooled α’β’ KCs and pooled γ KCs. Distances between food-related odour representations are shown in orange, between reproduction-related odour representations in blue, between danger-related odours in grey, and between food and other odour representations in brown. Significance of clustering of odour representations based on ethological groups was assessed using PERMANOVA (blue *, αβ KCs: p = 0.019; α’β’ KCs: p = 0.016; γ KCs: p = 0.455). The ability of KC types to segregate odour representations was examined by comparing distances within and between ethology-based groups (black *). αβ KCs displayed much greater distances between food-related odour representations when compared to between other odour representations (Kruskal-Wallis H-test, p < 0.001). However, γ KCs did not show significant differences in these comparisons (p = 0.16). Across KC types, αβ and α’β’ KCs displayed larger distances between food-related odours than γ KCs (Kruskal-Wallis H-test, αβ KC: p < 0.001, α’β’ KC: p = 0.01). * p<0.05, ** p<0.01, *** p<0.001. Error bar = Mean ± STD. (c) Comparison between response overlap and Euclidean distance of odour representations in αβ KC. The representations of food-related odours (orange) were farther apart, while the representations of reproduction-related odours (blue) were closer, even though both groups shared a similar response overlap. The representations of danger-related odours are labelled in grey. (d) Histogram of response strengths (average ΔF/F_0_) of αβ KCs to food-related (orange), reproduction-related (blue) or danger-related (grey) odours. The food-related odours more often elicit stronger responses than the other two categories. (e) Fraction of KCs responding to food-related odours per KC type. The colours of the stacked bar denote the number of food-related odours that the KCs responded to (1 - 4). A larger fraction of α’β’ KCs responded to food-related odours (77 %), followed by αβ KCs (50 %) and γ KCs (27 %). Specifically, 27.8 % of α’β’, 8.3 % of αβ and 6.8 % of γ KCs responded to all four food-related odours. (f) Euclidean distances between odour representations for the pooled all-KCs fly and the surrogate all-KCs fly. Distances between food-related odour representations are shown in orange, between reproduction-related odour representations in blue, between danger-related odours in grey, and between food and other odour representations in brown. Comparison between distances among food-related odour representations and distances between food-related and other odour representations (PERMANOVA, blue *; all-KCs fly: p = 0.01; surrogate all-KCs fly: p = 0.04). In addition, in both pooled all-KCs fly and surrogate all-KCs fly, distances between food-related odours were higher than that between other odours (black *, Kruskal-Wallis H-test, all-KCs fly: p < 0.001, surrogate all-KCs fly: p < 0.001). Error bar = Mean ± STD.

To evaluate grouping of odour representations based on ethological relevance into food-related and other (reproduction- and danger-related) odours, we applied multivariate ANOVA (PERMANOVA ^39^) to the Euclidean distances between odour representations for each KC type. By comparing the distances between the food-related odour representations, between the other odour representations, and between food-related and other odour representations, we found that αβ and α’β’ KCs are better at segregating food-related odours from the other odours than γ KCs (Fig. 3b). Thus, supporting the view that αβ and α’β’ KCs can represent food-related odours as a category.

The ability of a circuit to categorise related inputs might compete with its capacity to discriminate among those inputs ^40^. We therefore addressed the ability of KC types to discriminate between odours within the odour categories by comparing Euclidean (Fig. 3b) or cosine distances (Supplementary Figs. 3 - 6) within and between KC representations of ethology-based odour categories. Although food-related odours are physio-chemically more similar (Supplementary Fig. 2, ^41^), representations by αβ KCs displayed greater Euclidean distances between food-related odours when compared to reproduction-related or danger-related odours, highlighting potentially an enhanced capability to discriminate particularly between food-related odours (Fig. 3b). Furthermore, γ KCs did not show significant differences when comparing distances between food-related odours and between danger-related odours. In all KC types, the representations of reproduction-related odours were more clustered than the other odour categories (Fig. 3a, 3b), suggesting lower discrimination capacity. Across KC types, αβ and also α’β’ KCs displayed larger distances between food-related odours than γ KCs. These findings suggest that αβ and to a lesser extent also α’β’ KCs form divergent representations of food-related odours. Notably, although γ KCs have a connection bias towards RPNs, we did not observe increased distances between reproduction-related odour representations.

The overlap between cells responding to different odours is often used as a measure of discrimination (^21,42^, see Methods). A higher overlap is suggested to indicate more similar responses and less discrimination ^21,43^. Interestingly, αβ and α’β’ KCs exhibited similar overlap between cells responding to different food-related odours and between cells responding to different reproduction-related odours (Fig. 3c). Thus, the overlap level per se does not seem to be sufficient to explain the larger distances between representations of food-related odours in αβ KCs. Interestingly, we also did not see clear differences in the cosine distances between odour representations (Supplementary Figs. 3 - 6). However, larger proportions of αβ and α’β’ KCs responded to food-related odours with a higher dynamic range of responses, when compared to γ KCs (Fig. 3d and e). We suggest that these factors contributed to a larger effective volume of the activity space occupied by food-related odour-representations, and hence resulted in larger distances between food-related odour representations, contrary to what might be expected from response overlap alone.

Lastly, to ensure the analysed sample of KCs was representative, we created a "surrogate all-KCs fly" by pooling experimental data to form 290 αβ, 122 α’β’, and 214 γ KCs, matching the ratio of KC types in the Hemibrain dataset. The relationships between distances among representations of odour categories observed in the surrogate fly were consistent with those in all-KC flies (*OK107-Gal4*, n = 3), indicating that our type sampling accurately reflects the general KC response pattern (Fig. 3f).

Taken together, our findings show that αβ, and to a lesser extent also α’β’ KCs can support odour categorisation based on ethological relevance, segregating food-related odours within an activity space. In addition, αβ KCs form highly divergent representations of food-related odours that might lead to a greater discrimination capacity among them. Conversely, γ KCs did not exhibit significant segregation of odour categories and did not display discrimination capacity biased towards specific odour categories.

### Impact of PN-to-KC connection bias on KC type discrimination and categorisation capabilities

To explore how different connection logics affect the categorisation and discrimination capabilities of KC types, we developed network models with varying PN-to-KC connection logic. We then examined the simulated odour representations by comparing Euclidean distances and assessing the performance of linear classifiers based on these representations.

Our networks consist of OSN, PN, and KC layers (Fig. 4a). The OSN and PN layers are based on previous models of the *Drosophila* antennal lobe ^44,45^, containing 50 OSN glomeruli that excite 103 PNs of 50 different types. Lateral feedforward inhibition is provided to the PNs by a local inhibitory neuron (LN). The odour responses of OSNs were taken from the extensive experimental data ^33,36,46,47^. The KC layer consists of three KC types: 799 αβ, 335 α’β’, and 590 γ neurons. Each KC is modelled as a rectified linear unit, that sums inputs from PNs along with feedforward inhibition from the APL neuron ^22^, which was kept global for simplicity. We explored three different PN-to-KC connection models: the *Hemibrain,* the *random uniform* and the *random non-uniform* (Fig. 1e, Methods). For each KC type we adjusted three parameters: the spiking threshold, the excitability, and the strength of inhibitory input from the APL neuron. We optimized these parameters such that the lengths of the odour representation vectors for the Hemibrain connection model best fit those that we observed experimentally (see Methods). The spiking threshold was highest for γ, followed by αβ and then α’β’ KCs. α’β’ KCs exhibited the highest excitability, followed by αβ and then γ KCs. The strengths of inhibitory input were similar for the three KC types. This is in agreement with reported electrophysiological properties of KC types ^48^.

**Figure 4.**
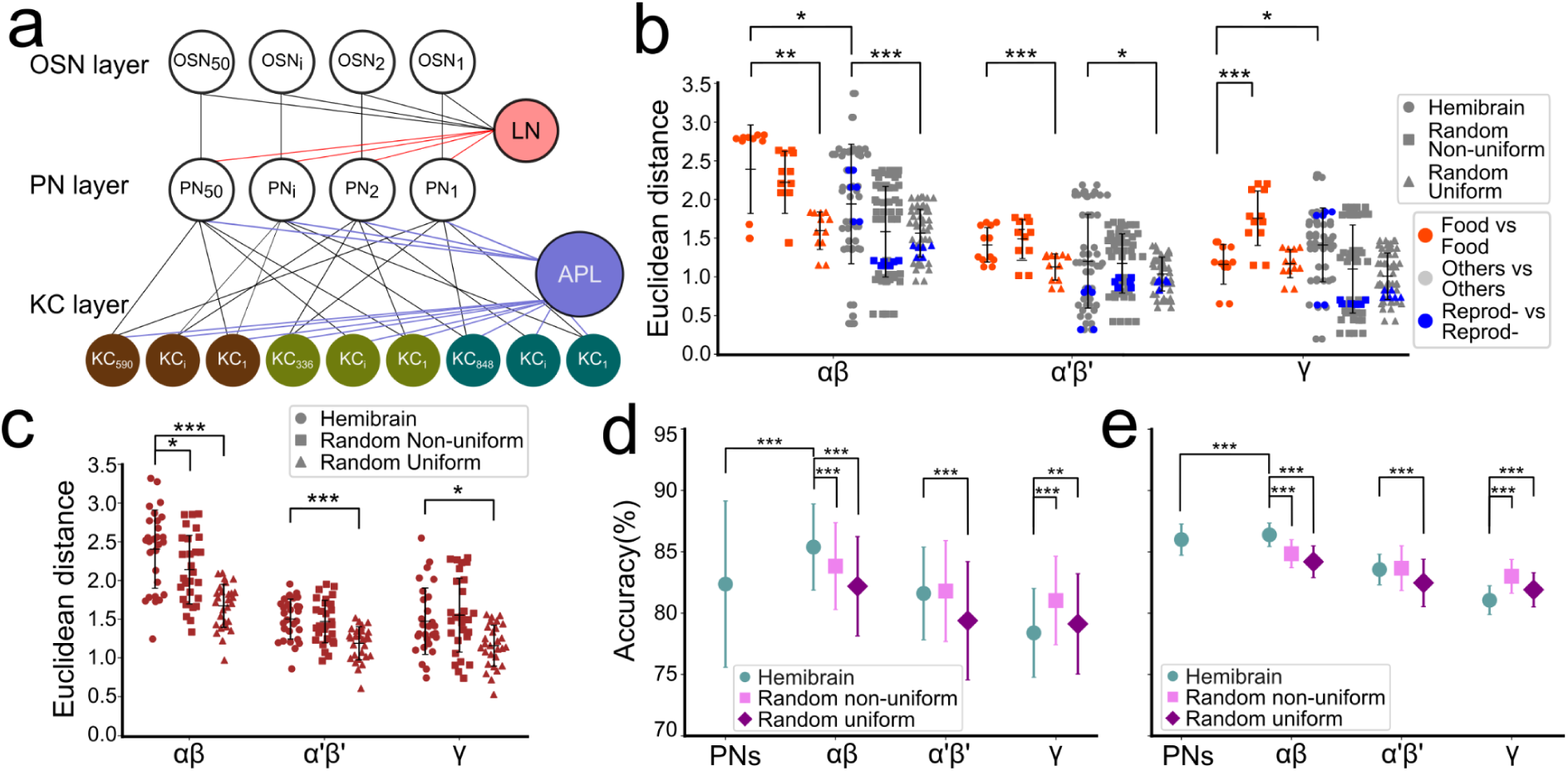
Network modelling of odour representations of different KC types using Hemibrain and random connection models. (a) The model consists of an OSN layer, a PN layer, and a KC layer. 50 OSN glomeruli activate 103 PNs across 50 PN types. The KCs layer linearly sums PN layer activity. There are three types of KCs: 799 αβ KCs (dark green), 335 α’β’ KCs (light green), and 590 γ KCs (brown). The LN and the APL neuron provide feedforward inhibition. (b) Euclidean distances between the KC odour representations for various odour categories, comparing responses in the Hemibrain (circles), random non-uniform (square), and random uniform connection models (triangles). Distances between food-related odours are shown in orange, between reproduction-related odours in blue, and between other odours in grey. * p<0.05, ** p<0.01, *** p<0.001, Kruskal-Wallis H-test. Error bar = Mean ± STD. (c) Same as in (b) but between food-related and other odour representations. (d) Linear classifier accuracy for discrimination between food-related odours for the different cell types in the three connection models: Hemibrain (circles), random non-uniform (squares), and random uniform (diamonds). * p<0.05, ** p<0.01, *** p<0.001, Kruskal-Wallis H-test. Error bar = Mean ± STD. (e) Same as in (d) but for categorisation into food-related and other odours.

For αβ and α’β’ KCs, the Hemibrain and the random non-uniform connection models had higher capacity to separate food-related odour representations than the random uniform model, in terms of distances between odour representations (Fig. 4b). Conversely, for γ KCs, the Hemibrain and the random uniform connection models exhibited similar distances between food-related odour representations, both shorter than the random non-uniform model (Fig. 4b). On the other hand, for αβ KCs, the segregation of food-related odour from other odour representations was most pronounced in the Hemibrain connection model, and least pronounced in the random uniform connection model for all KC types (Fig. 4c). Thus, the different PN-to-KC connection logics have a clear impact on the distances between odour representations.

Considering the larger distances between food-related odour representations in αβ KCs compared to the other KC types observed experimentally (Fig. 3b), as well as in the simulations (Fig. 4b), we hypothesized that αβ KCs would support superior discrimination between food-related odours. To thus assess discrimination performance, we randomly assigned positive or negative valences to food-related odours. We then trained linear classifiers to distinguish the valence based on odour representations of each KC type or of PNs. Using the Hemibrain connection model, the linear classifiers trained with αβ KCs representations displayed the highest discrimination accuracy among food-related odours, followed by the classifiers trained on PNs, α’β’ and then γ KCs representations (Fig. 4d). When comparing the different connection models, for both αβ and α’β’ KCs the Hemibrain model led to a better discrimination performance than the random uniform model. On the other hand, it resulted in a more similar performance to the random non-uniform connection model (Fig. 4d). Conversely, the Hemibrain connection model led to the worst performance of a γ KCs representations-trained classifier (Fig. 4d). These findings suggest that the random non-uniform connections could account to a certain extent for the enhanced food-related odour discrimination performance observed for αβ or α’β’ KCs. However, the Hemibrain connections do not seem to endow γ KCs with representations that support better food-related odour discrimination.

Given the larger distances between representations of food-related and other odours observed experimentally (Fig. 3b), as well as in the simulations (Fig. 4c), we hypothesized that αβ KCs would support superior categorisation between food-related and other odours. To assess the model’s categorisation performance, we trained linear classifiers to distinguish between food-related and other odours based on the representations formed by PNs and each KC type. Indeed, in the Hemibrain connection model, αβ KC representations supported the highest categorisation accuracy, followed by PN, α’β’ and γ KC representations (Fig. 4e). When comparing the different connection models, the Hemibrain model outperformed the others for αβ and for α’β’ KC representations-trained classifiers (Fig. 4e). Meanwhile, γ KC representations in the Hemibrain connection model led to the worst performance in this categorisation task than either random model (Fig. 4e). These results indicate that while Hemibrain connections support odour categorisation in αβ and α’β’ KC representations, they appear to diminish it for γ KC representations.

In summary, the random non-uniform connections seem to produce αβ and α’β’ KC representations that support enhanced food-related odour discrimination as well as categorisation into food-related and other odours. Nevertheless, in αβ KCs, the Hemibrain connections contribute to the highest categorisation and discrimination performances, potentially due to oversampling of FPNs (Fig. 1F). Meanwhile, γ KCs, with their connections resembling the random uniform model, adopt a more generalist approach.

## Discussion

Our findings offer a new perspective on how the MB calyx organisation defines the olfactory representations in KCs. By combining the analysis of connectome data with functional recordings of KCs responses and computational modeling of the circuit, we find that the three main KC types form odour representations with distinct properties: odour representations formed by αβ and α’β’ but not γ KCs efficiently support the categorisation of food-related odours from other odours, as well as discrimination among the food-related odours. This can be largely attributed to the specific patterns of PN-to-KC connections realised by KC types.

Using data from the Hemibrain dataset, we identified a significant connection bias of the αβ and α’β’ KCs towards FPNs, corroborating and extending the findings based on the FAFB dataset ^20^. In contrast, γ KCs display a connection pattern that more closely resembles random sampling. We note here that these data derive from adult female brains, it is so far unclear how these biases might apply to the male brains. Through our analysis, we show that distinct connection logics coexist in the complex calycal circuit, including biased connections that diverge from the randomness often hypothesised for cerebellum-like circuits ^49,15,30^. Recent findings suggest how PN-to-KC connection biases in the calyx might emerge during development. The boutons of some PN types are distributed in the calyx in a reproducible manner, while KC dendritic claw positioning depends on both KC neuroblast origin and KC type ^50–52^. In addition, αβ and α’β’ KCs claws wrap the FPNs’ boutons within microglomeruli more than boutons of other PN types ^20^. Complementing developmentally defined programs, the olfactory environment might modulate the PN-to-KC connections. In fact, the number of boutons formed by a PN depends upon the neuron’s activation in the first days after eclosion ^53–55^. Nonetheless, partially anosmic flies seem to display unchanged PN-to-KC connection patterns ^56^. Further work is thus needed to unravel how distinct connection logics—biased and random—are established and refined during development and how they might be related to evolutionary constraints ^57^.

We discovered that αβ and α’β’ KCs segregate odour representations based on the ethological relevance of the odours, particularly food-related odours from other odours, supporting odour categorisation. We propose that grouping various odours under a unified ‘food’ category may simplify decision-making and enhance behavioural response efficiency. This type of processing seems in conflict with the need to isolate distinctive aspects among odours for their discrimination ^40^. Nonetheless, αβ and α’β’ KCs representations support both operations and, while grouping food-related odours in a category, they keep the odours in this category separated. Discriminating nuances among food-related odours, such as differentiating ripeness, could enable flies to select optimally nutritious food sources.

Importantly, specific odours within the food category could be individually associated with experiences, allowing the animals to carry out efficient foraging ^1–3^. Our modelling data indicate that the capacity of αβ and α’β’ KCs to separate food-related odour representations can be largely explained by the random non-uniform connections. Thus, biases in the number of boutons formed and the number of connections per bouton might be sufficient to support food-related odour discrimination by αβ or α’β’ KCs.

γ KCs form their connections with PNs following a different logic than αβ or α’β’ KCs and serve a distinct role in olfactory processing. γ KCs have more claws and a higher activation threshold compared to αβ and α’β’ KCs, and tend to respond to a single or very few odours^48^. We show here that γ KCs form less biased connections with PN categories and do not differentiate between categories of odours, at least for the set of odours we tested. Their responses can be reproduced by a random uniform connection model. Taken together, it appears that γ KCs fulfil a role originally proposed for the MB—forming distinct response patterns to odours irrespective of their type. Such versatility may allow animals to flexibly associate value with new odours based on experiences, which may be advantageous for adaptive responses in a dynamic environment.

Along these lines, a recent study suggested that odour representations in KCs generally align more closely with the natural source space than with the PNs chemical descriptor space ^58^. This supports a strategic reorganisation of odour coding at the KC level. In analogy to the *Drosophila* MB, neurons in the rodent piriform cortex randomly integrate inputs from multiple glomeruli ^59^ and reconfigure odour representations they receive from the olfactory bulb, emphasizing relationships among odours ^60^.

Different KC types were found to support various stages of memory formation or retrieval ^24–29^. In this context, short-term memory is primarily supported by α’β’ and γ KCs, while long-term memory relies on αβ and γ KCs ^61^. How the role of different KC types in short-term or long-term memory intersects with their specific responses toward different odour categories reported here requires future investigation. Nonetheless, emerging evidence suggests specialised roles for specific KC types in behaviours related to particular odours. For instance, αβ KCs are crucial for the persistent tracking of food-related odours in the absence of immediate rewards in hungry flies ^62^. γ KCs, on the other hand, are central to courtship conditioning in male flies, associating the reproductive pheromone cVA with aversive experiences ^63^. In light of our findings, it is plausible that specialised roles of αβ and γ KCs may arise from their connection preferences towards FPNs and RPNs, respectively.

In the rodent cerebellar circuit, connections from mossy fibers to the granule cells were recently shown to be partially biased ^64^, akin to the biased sampling of FPNs by αβ and α’β’ KCs. Computational modeling suggested that this bias in the cerebellar circuitry can lead to higher representation dimensionality and better pattern separation performance. In comparison to classic random circuit models this also improved resilience to noise for the biased inputs ^64^. Considering the conserved organisation of cerebellum-like circuits, including these recently emerging parallels, our data will help refine the current models of this canonical circuit motif that supports learning.

## Acknowledgements

We thank the Light Microscopy Facility at DZNE and Luigi Prisco for their support in technical development and Phuong Tran, Rita Kerpen, Magdalena Magiera, and Olga Sharma for technical assistance. We are grateful to the Bloomington Stock Center, Kyoto Stock Center, and Lisa Scheunemann for providing fly lines. We also thank Andrew Lin, Hokto Kazama, Tobias Ackels, and Lisa Scheunemann for critical discussions and feedback on the manuscript. Special thanks to the members of the Tavosanis Lab - Karolína Doubková, Jonathan Smart, Pallavi Syam, and Gokul Madhav for their critical reading of the manuscript. We are also grateful to Raoul-Martin Memmesheimer and Dan Ehninger for their support during the project. This work was supported by the DFG FOR 2705 grant and the “iBehave network” (funded by the Ministry of Culture and Science of the State of North Rhine-Westphalia, Germany) to G.T..

## Author Contributions

Ivy Chi Wai Chan: Conceptualization, Experiments, Data analysis, Modelling, Project administration, Writing of original draft, Review and editing;

Felipe Yaroslav Kalle Kossio: Data analysis, Modelling, Review and editing;

Gaia Tavosanis: Conceptualization, Funding acquisition, Project administration, Supervision, Review and editing.

## Material and Methods

| Experimental Models: Organisms/Strains |  |  |
| --- | --- | --- |
| D. Melanogaster:<br>*OK107-Gal4* | Bloomington Drosophila<br>Stock Center | Bloomington ID:854 ;<br>FlyBase ID: FBti0004170 |
| D. Melanogaster:<br>*c305a-Gal4* | Bloomington Drosophila<br>Stock Center<br><sup>65</sup> | Bloomington ID: 30829 ;<br>FlyBase ID: FBal0215258 |
| D. Melanogaster:<br>*GMR71G10-Gal4* | Bloomington Drosophila<br>Stock Center<br><sup>66</sup> | Bloomington ID: 39604;<br>FlyBase ID: FBtp0062423 |
| D. Melanogaster:<br>*NP3061-Gal4* | Kyoto Stock Center<br><sup>67</sup> | DGRC Number: 104360<br>FlyBase ID:FBti0035130 |
| D. Melanogaster:<br>*UAS-nls-mCherry,*<br>*UAS-GCaMP6f* | Gift from L. Scheunemann<br><sup>68</sup> | FlyBase ID: FBal0281812,<br>FBtp0071546 |
| Chemical Reagents: |  |  |
| Odourants | CAS number, company<br>and catalogue number | Final dilution |
| 1-hexanol (Hex) | CAS #: 111-27-3<br>Sigma-Aldrich<br>471402 | 1:100 |
| 2-nonanone (Non) | CAS #: 821-55-6<br>Th. Geyer<br>11812319 | 1:100 |
| Acetone (Ace) | CAS #: 67-64-1<br>Th. Geyer<br>11647213 | 1:100 |
| Butyl butyrate (BB) | CAS #: 109-21-7<br>VMR International<br>B24390.AK | 1:100 |
| 11-*cis*-Vaccenyl acetate<br>(cVA) | CAS #: 6186-98-7<br>Pherobank | 1:100 |
|  | 10421 |  |
| Ethyl acetate (EA) | CAS #: 141-78-6<br>Th. Geyer<br>11835215 | 1:100 |
| Ethyl hexanoate (EH) | CAS #: 123-66-0<br>Santa Cruz<br>sc-235049 | 1:100 |
| Farnesol (Fa) | CAS #: 106-28-5<br>MP Biomedicals<br>0210168005 | 1:100 |
| Methyl laurate (ML) | CAS #: 111-82-0<br>Sigma-Aldrich<br>07041-100MG | 1:100 |
| Methyl palmitate (MP) | CAS #: 112-39-0<br>Sigma-Aldrich<br>P5177-5G | 1:100 |
| Pentyl acetate (PA) | CAS #: 628-63-7<br>Sigma-Aldrich<br>46022-250ML-F | 1:100 |
| Pyrrolidine (Pyr) | CAS #: 123-75-1<br>Sigma-Aldrich<br>P73803-100ML | 1:1000 |
| Mineral oil | CAS #: 8042-47-5<br>Sigma-Aldrich<br>330779-1L |  |

### Fly husbandry

Flies were raised in a 12h/12h light-dark cycle on a standard cornmeal-based diet at 25-27°C, 60 % relative humidity. All functional imaging experiments were performed on adult female flies 3–5 days after eclosion. To drive the expression of GCaMP6f and NLS-mCherry in different KC types, the following Gal4 drivers were utilised: *OK107-Gal4* for all KC types, *NP3061-Gal4* for αβ KCs, *c305a-Gal4* for α’β’ KCs, and *GMR71G10-Gal4* for γ KCs. We used FlyBase (release FB2024_06) to find information on the available stock lines^69^ (for further detail, see Experimental Models: Organisms/Strains).

### Global mapping of PN clusters from FAFB and Hemibrain

PN types were grouped based on their ethological roles following ^20^. Assignment of D adPNs and DC3 adPNs deviates from ^20^, which was motivated by ^32,70^.

**Table 2.** Ethology-based grouping of PN types.

|  |  |
| --- | --- |
| Food-odour-responding PNs (FPNs) | DL2v adPNs, DM1 IPNs, DM2 IPNs, DM3 adPNs, DM4 adPNs, DP1m adPNs, VA4 IPNs, VM2 adPNs, VM3 adPNs |
| Reproduction-odour-responding PNs (RPNs) | D adPNs, DA1 IPNs, DC1 adPNs, DC3 adPNs, DL3 IPNs, VA1d adPNs, VA1v adPNs, VC5 adPNs |

The global mapping of PN clusters was conducted using the FAFBv14 and Hemibrain:v1.1 datasets. Individual PNs were identified by their accession numbers in both datasets. These neuron objects were then retrieved using the VFB CATMAID server for FAFB and the NAVis-neuprint interface for Hemibrain. To construct a global map, PNs from the FAFB dataset (encompassing both right and left hemispheres) were aligned onto the Hemibrain template (right hemisphere) alongside the native Hemibrain PNs.

Specifically, PNs from the left hemisphere in the FAFB dataset were mirrored to the right hemisphere using NAVis’s ‘mirror brain’ function. Subsequently, both the mirrored (left hemisphere) and original (right hemisphere) FAFB PNs, as positioned in the FAFB14 template, were transformed to the Hemibrain template (JRCFIB2018Fraw, right hemisphere) employing the ‘xform brain’ function. Hemibrain PNs were plotted on the same Hemibrain template (JRCFIB2018Fraw), together with the transformed FAFB PNs.

To enhance the visualisation of PN distribution within the calyx, PN neuropils were pruned according to the Hemibrain calyx volume using the ‘prune by volume’ function. Additionally, synaptic locations for the selected PNs were extracted and mapped onto the Hemibrain template for analysis of PN synapse distributions.

### Wasserstein distance between PN presynapses and KC postsynapses

We obtained locations of PN presynapses and KC postsynapses within the calyx from the FAFB dataset. We constructed pre- and postsynapse distributions by dividing calyx into voxels measuring 10 x 10 x 10 µm³ and counting the numbers of pre- or postsynapes within each voxel. To quantify the distance between the distributions of PN postsynapses and KC presynapses, we employed the Wasserstein metric ^71^. This symmetric metric is the minimum “cost” needed to transform one distribution into the other, effectively capturing the similarity between two probability distributions. In our case, cost is the number of pre- or postsynapses moved times the average distance they were moved.

### Analysis of the Hemibrain PNs-to-KC type connections

We analysed the connections from each PN type to each KC type i.e. αβ, α’β’ and γ KCs using data from Hemibrain:v1.1. In this analysis, we only considered connections with a weight exceeding five active zones. To create a KC type-specific weighted connection matrix, individual PNs were categorised based on their type or associated glomerulus. Columns representing the same PN type within the matrix were then summed to form a single column per PN type, thus compiling the Hemibrain PNs-to-KC type connection matrix. This matrix was further analysed to quantify the proportion of inputs from various PN groups—such as FPNs, RPNs, and others—to each KC type. This was achieved by summing the connection weights for each group, providing a detailed measure of the synaptic contributions to the KCs from different PN types.

### Random PNs-to-KCs connection models

To simulate random connections between PNs and KCs, we developed two models to generate PN-to-KC connectivity matrices. Each KC’s number of dendritic claws and the connection weights (i.e. number of synapses per claw) were drawn from the distribution specific to each KC subtype in the Hemibrain: αβ KCs had an average of 5 claws, α’β’ KCs 4, and γ KCs 8; with connection weights of 14 ± 8 for αβ and α’β’ KCs, and 18 ± 8 for γ KCs. In the *random uniform connection model*, each KC claw had an equal chance of connecting to one of the 50 PN types. In the r*andom non-uniform* connections model, each KC claw was connected to a PN type with probability proportional to the actual fraction of connections formed by that PN type in the Hemibrain dataset. Connection weights were drawn from the distribution specific to each KC type in the Hemibrain.

### *In*-*vivo* calcium imaging

For each imaging experiment, female adult flies of 3-6 days post-eclosion were briefly anaesthetised on ice, and positioned in a polycarbonate imaging chamber. To allow optical access to the KCs cell bodies, a small window was opened through the head capsule under Ringer’s solution (5 mM HEPES, pH 7.4, 130 mM NaCl, 5 mM KCl, 2 mM CaCl_2_, 2 mM MgCl_2_; pH adjusted to 7.2). To minimise movement, fly heads were stabilised with 1.2 % low melting agarose (Thermo Scientific) in Ringer’s solution, immediately before dissection. Flies were imaged with a two-photon laser-scanning microscope (LaVision BioTec, TriM Scope II) equipped with an ultra-fast z-motor (PIFOC Objective Scanner Systems 100 μm range) and a Nikon 25× CFI APO LWD Objective, 1.1 NA water-immersion objective.

Both GCaMP6f and NLS-mCherry were excited at 1,000 nm. Bandpass filters for 525/550 nm and 595/640 nm were used for detecting the GCaMP and mCherry fluorescence signals, respectively. For volumetric imaging to image most of the KCs cell bodies (a volume of 100 µm x 100 µm x 25 µm), a 3D stack of 6 z-sections with 2-3 µm intervals was acquired at around 1 Hz.

### Odours stimulation protocol

Individual monomolecular odours were used to stimulate the olfactory responses of KCs. Odour stimuli were delivered to the fly using a 220A olfactometer (Aurora Scientific). The odours identity and dilutions are listed in the *Reagents section* above. Upon delivery to the fly, the olfactometer further diluted the odour 1:10 in air. Each trial consisted of a series of 8 to 12 odours presented in a pseudo-random order for 2 seconds each with a 45-second interval between presentations. For each fly, we carried out 2 to 4 trials, allowing a 10-minute rest interval between them.

### Image registration

For each fly, all imaging trials were compiled into a unified dataset. We utilised the cytoplasmic GCaMP6f to monitor odour-evoked calcium activity and the nuclear NLS-mCherry for structural labelling, essential for accurate motion correction and cell segmentation. Our volumetric imaging comprised stacks of 6 planes per frame. Frames were registered using the Advanced Normalisation Tools (ANTs) ^72^.

We manually identified 6-7 landmark KCs per fly and tracked these through truncated time series, which included 20 frames before and 9 frames after odour onset for all odours. The positions of other KCs were identified from an average image and predicted across the frames using thin plate spline transformations.

To extract GCaMP6f signals, circular masks of 5 µm diameter were created based on the ROI locations. Due to the dense packing of KCs, which could lead to signal contamination from adjacent cells, overlapping pixels were selectively excluded.

For each odour presented, the baseline fluorescence (F_0_) was established using the 20 frames preceding the odour onset. The normalised response of a KC for the corresponding odour at time t was calculated by (F_t_ - F_0_)/ F_0_, where F_t_ represents the fluorescence intensity at time t. The odour-evoked average activity for each KC was calculated by averaging across all sessions and across time (from odour onset to 5 seconds post-onset). The analysis included all KCs, regardless of their level of activity. All KCs responses from flies of the same genotype were pooled together.

### Similarity analysis of the functional responses of KC types

To investigate the structural organisation of olfactory representations across different KC types, for each type we constructed odour representation vectors from the odour-evoked average activities of all KCs of the same type.

We quantitatively assessed the discriminative ability of each KC type by calculating Euclidean distances and correlation distances within and between groups of odour representation vectors. Euclidean Distance calculates the straight-line distance between two points in N-dimensional space and is defined as:

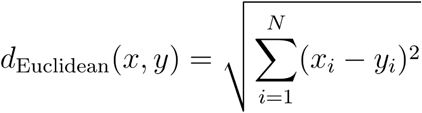

where *x_i_* and *y_i_* are the components of the vectors *x* and *y*, respectively.

Similarly, the cosine distance between two odour representation vectors measures the angular distance between the two vectors. It is defined as:

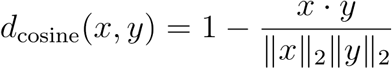

where and *x* · *y* is the dot product of the vectors *x* and *y*.

Intra-group distances were measured between vectors within the same ethological odour category, while inter-group distances were measured between vectors from different categories. By comparing these distances, we could quantify how distinctly each KC type represents odours that belong to different ethological categories.

Additionally, we analysed the overlap between cells that responded to multiple odours within each KC type, as a higher overlap can suggest more similar responses and less discrimination. To determine if a KC responded to a given odour, we applied a threshold of 4 × STD above the baseline fluorescence (F_0_) to the odour-evoked average activities. Overlap was calculated for pairs of odours, *A* and *B*, as follows:

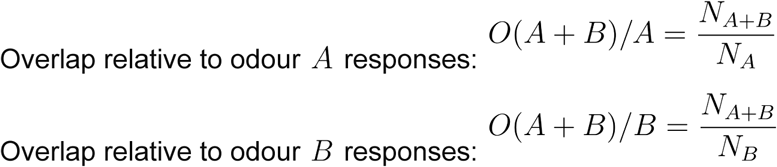

, where *N_A_* denotes the number of KC responding to odour *A, N_B_* denotes the number of KC responding to odour *A*, and *N_A+B_* denotes the number of KC responding to both odours *A* and *B*.

Both values, *O*(*A* + *B*)/*A* and *O*(*A* + *B*)/*B*, are considered to account for the different numbers of responding KCs per odour. This approach enables a balanced evaluation of response overlap.

### Statistical analysis

To assess the significance of the groupings of KC representations according to the ethological functions of the odours, we employed Permutational Multivariate Analysis of Variance (PERMANOVA). This non-parametric statistical test compares the differences across multiple groups based on their multivariate data distributions. Unlike traditional ANOVA, PERMANOVA utilises a distance matrix—specifically, the Euclidean distance matrix in our study—to determine if the observed grouping in KC representations is statistically significant.

### Network model of the MB input circuit

Our model of the *Drosophila* MB input circuit integrates the primary olfactory sensory neurons (OSNs) and the secondary layer of olfactory projection neurons (PNs), drawing inspiration from existing models of the *Drosophila* antennal lobe ^44,45^. The first layer contains 50 OSNs corresponding to 50 antennal lobe glomeruli, which transmit signals to 103 PNs, of 50 PN types, through feedforward excitation. The OSNs’ activities were based on experimental data sourced from various studies ^33,36,46,47^, with a local inhibitory neuron (LN) providing lateral feedforward inhibition. The response of each PN to specific odours, *r*_PN_, was modelled by the following equation:

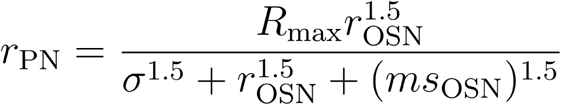

Here, *r*_OSN_ represents the firing frequency of the OSN in response to an odour, *σ* is a model fitting constant, mmm denotes the inhibition sensitivity of the glomeruli, and *R*_max_ is the maximum response rate of the PNs, capped at 165 Hz. Given that the available datasets typically cover only 28 glomeruli, responses for the remaining 22 glomeruli are set to be zero.

The KC layer is composed of three KC types: 799 αβ, 335 α’β’, and 590 γ neurons. Each KC sums inputs from the PNs and integrates feedforward inhibition from the APL neuron ^22^, maintained globally for simplicity:

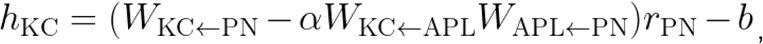

where *W*_KC←PN_ denotes the number of active zones from each PN bouton to each KC claw, *α* is the inhibition gain for the APL neuron, *W*_*KC*←APL_ and *W*_APL←PN_ represent the number of active zones from the APL neuron to KCs and that from PNs to the APL neuron respectively, and *b* is the KC’s response threshold. *W*_KC←PN_ were generated based on the aforementioned connection models (See above: Analysis of the Hemibrain PNs-to-KC type Connection and Random PNs-to-KCs Connection Models). *W*_APL←PN_ were randomly sampled from a log-normal distribution reflective of Hemibrain data (mean ± STD = 3.79 ± 0.48), while *W*_*KC*←APL_ were drawn from a normal distribution, specific to each KC type in the MB calyx (αβ KC: 9.357 ± 3.924; α’β’ KC: 9.191 ± 4.309; γ KC: 23.366 ± 4.986).

To compute the responses of KCs, we employed a Rectified Linear Unit (ReLU) function:

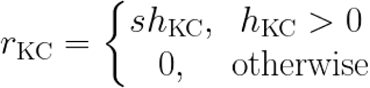

Here, *s* signifies the slope of the activation function, modelling the activation dynamics of the KCs as a function of the total input *h_KC_*.

For each KC type, we adjusted three parameters - spiking threshold (*b*), the strength of inhibitory input from the APL neuron (*α*), and the excitability (*s*). Using the Hemibrain connection model, we optimised these parameters using the Nelder-Mead algorithm such that lengths of resulting odour representation vectors best fitted the experimentally observed ones from our odour panel.

### Linear classifiers of the network model

Four additional food odours (acetic acid, 2,3-butanedione, ethyl lactate, and hexyl acetate) and one additional non-food odour (geosmin) were added to the experimental odour panel (Table 3), forming a set of 8 food and 8 non-food odours for linear classification. Gaussian noise with a standard deviation of 40 Hz was applied to the OSN activity to generate 100 noisy OSN odour representations. The OSN activity was clipped at zero to prevent negative values. Half of the generated odours were used to train the linear classifier, implemented as a support vector machine (SVM), while the remaining half were used to test the classifier’s accuracy. The classifiers were trained using the activity of different KC types either separately or combined.

**Table 3.** The panel of 4 additional food-related odours and 1 danger-related odour.

| Ethological relevance | Additional Odours | Primarily Activated Glomeruli | Reference |
| --- | --- | --- | --- |
| Food | Acetic acid | DL2, DP1 | <sup>32</sup> |
| Food | 2,3-butanedione | VA2 | <sup>32</sup> |
| Food | Ethyl lactate | VC4 | <sup>32</sup> |
| Food | Hexyl acetate | DM2 | <sup>32</sup> |
| Danger | Geosmin | DA2 | <sup>73</sup> |

To assess discrimination performance, half of the food odours were assigned a valence (positive or negative), and the classifier was trained to differentiate between these valances. This procedure was repeated 100 times for robustness. A similar approach was applied to the non-food odours.

For categorization testing, the classifiers were trained to distinguish between food and non-food odours, following the same methodology.

## Supplementary Material

**Supplementary Table 1.** Number of connections from each PN type to all KCs in the Hemibrain dataset. The number of connections (one PN bouton to one KC claw) each PN type forms with KCs. In the random non-uniform connection model, the probability that a KC claw connects to a given PN type is proportional to the actual fraction of connections that PN type forms in the Hemibrain dataset.

| PN type | Number of PNs | Number of connections | Connection probability | Connection probability per neuron |
| --- | --- | --- | --- | --- |
| D | 3 | 188 | 0.0177 | 0.0059 |
| DA1 | 7 | 334 | 0.0314 | 0.0045 |
| DA2 | 5 | 164 | 0.0154 | 0.0031 |
| DA3 | 2 | 44 | 0.0041 | 0.0021 |
| DA4l | 1 | 83 | 0.0078 | 0.0078 |
| DA4m | 1 | 27 | 0.0025 | 0.0025 |
| DC1 | 1 | 393 | 0.0369 | 0.0369 |
| DC2 | 2 | 262 | 0.0246 | 0.0123 |
| DC3 | 3 | 255 | 0.0240 | 0.0080 |
| DC4 | 1 | 63 | 0.0059 | 0.0059 |
| DL1 | 2 | 360 | 0.0338 | 0.0169 |
| DL2d | 5 | 208 | 0.0196 | 0.0039 |
| DL2v | 3 | 146 | 0.0137 | 0.0046 |
| DL3 | 1 | 59 | 0.0055 | 0.0055 |
| DL4 | 1 | 48 | 0.0045 | 0.0045 |
| DL5 | 1 | 173 | 0.0163 | 0.0163 |
| DM1 | 1 | 438 | 0.0412 | 0.0412 |
| DM2 | 2 | 372 | 0.0350 | 0.0175 |
| DM3 | 1 | 224 | 0.0211 | 0.0211 |
| DM4 | 1 | 327 | 0.0307 | 0.0307 |
| DM5 | 2 | 233 | 0.0219 | 0.0110 |
| DM6 | 4 | 348 | 0.0327 | 0.0082 |
| DP1l | 1 | 321 | 0.0302 | 0.0302 |
| DP1m | 1 | 521 | 0.0490 | 0.0490 |
| V | 2 | 225 | 0.0212 | 0.0106 |
| VA1d | 3 | 138 | 0.0130 | 0.0043 |
| VA1v | 3 | 97 | 0.0091 | 0.0030 |
| VA2 | 1 | 315 | 0.0296 | 0.0296 |
| VA3 | 2 | 270 | 0.0254 | 0.0127 |
| VA4 | 1 | 270 | 0.0254 | 0.0254 |
| VA5 | 3 | 188 | 0.0177 | 0.0059 |
| VA6 | 1 | 244 | 0.0229 | 0.0229 |
| VA7I | 1 | 135 | 0.0127 | 0.0127 |
| VC1 | 1 | 138 | 0.0130 | 0.0130 |
| VC2 | 1 | 176 | 0.0165 | 0.0165 |
| VC3I | 3 | 342 | 0.0321 | 0.0107 |
| VC3m | 3 | 289 | 0.0272 | 0.0091 |
| VC4 | 3 | 237 | 0.0223 | 0.0074 |
| VC5 | 1 | 151 | 0.0142 | 0.0142 |
| VL1 | 2 | 43 | 0.0040 | 0.0020 |
| VL2a | 1 | 223 | 0.0210 | 0.0210 |
| VL2p | 1 | 291 | 0.0274 | 0.0274 |
| VM1 | 1 | 73 | 0.0069 | 0.0069 |
| VM2 | 2 | 174 | 0.0164 | 0.0082 |
| VM3 | 2 | 181 | 0.0170 | 0.0085 |
| VM4 | 2 | 86 | 0.0081 | 0.0040 |
| VM5d | 5 | 343 | 0.0322 | 0.0064 |
| VM5v | 2 | 146 | 0.0137 | 0.0069 |
| VM7d | 2 | 147 | 0.0138 | 0.0069 |
| VM7v | 2 | 125 | 0.0118 | 0.0059 |
| **Total** | 103 | 10638 | 1.0 | 1.0 |

**Supplementary Figure 1.**
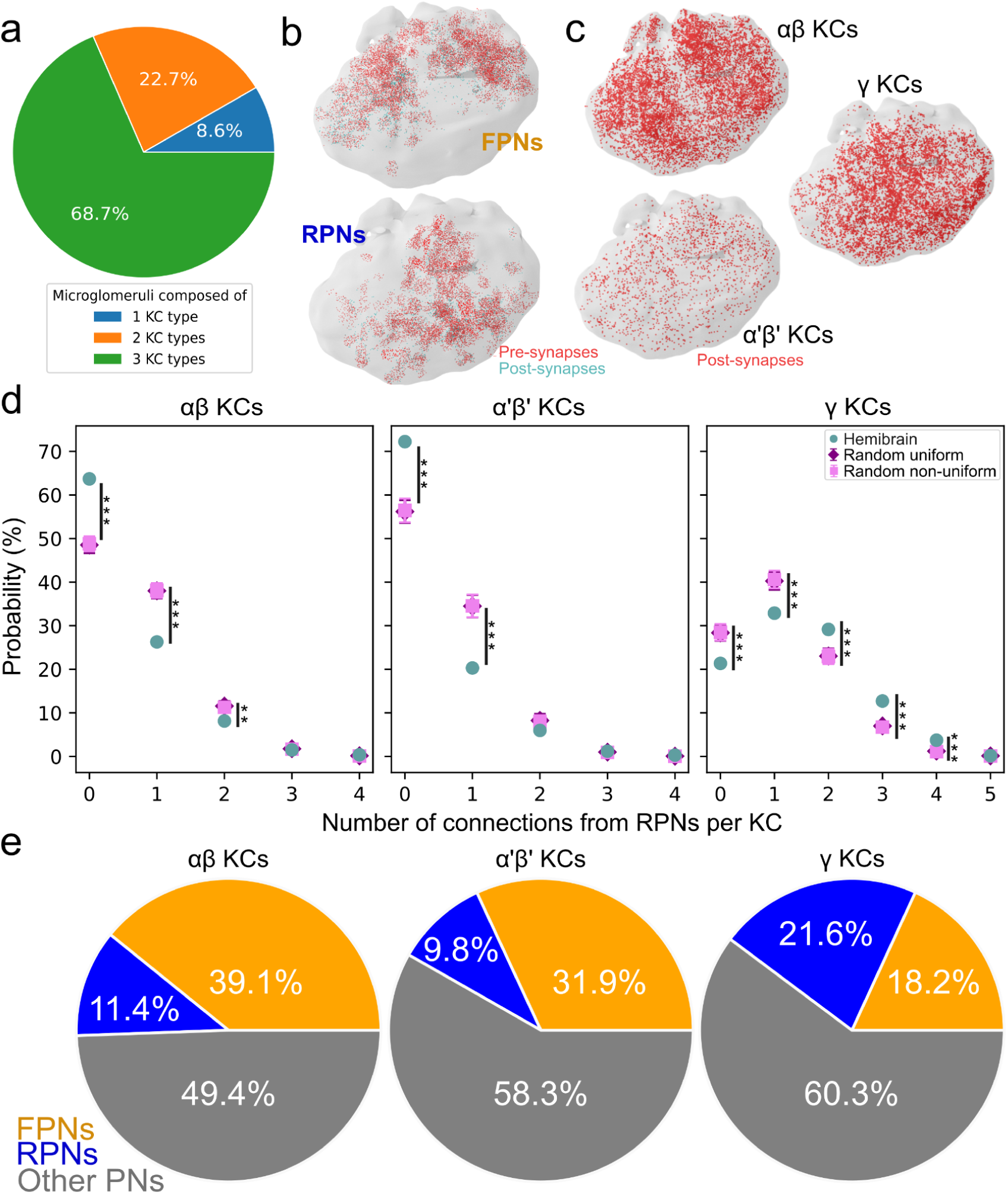
Synaptic distribution and PNs-to-KC connectivity in the MB calyx. **(a)** Number of KC types per microglomerulus (FAFB). **(b)** Locations of pre- and postsynapses of FPNs and RPNs in the MB calyx (Hemibrain and FAFB combined). **(c)** Locations of post-synapses of different KC types in the MB calyx (Hemibrain and FAFB combined). **(d)** KCs to RPNs connection probabilities in Hemibrain or in the two random connections models. The graph shows the probability that a KC forms a certain number of connections with RPNs in the Hemibrain (turquoise), in the random uniform (purple) and the random non-uniform (pink) connection models. * p<0.05, ** p<0.01, *** p<0.001, Kruskal-Wallis H-test. Error bar = Mean ± STD. **(e)** Proportion of number of connections from FPNs (orange), RPNs (blue) and other PNs (grey) to αβ KCs, α’β’ KCs or γ KCs.

**Supplementary Figure 2.**
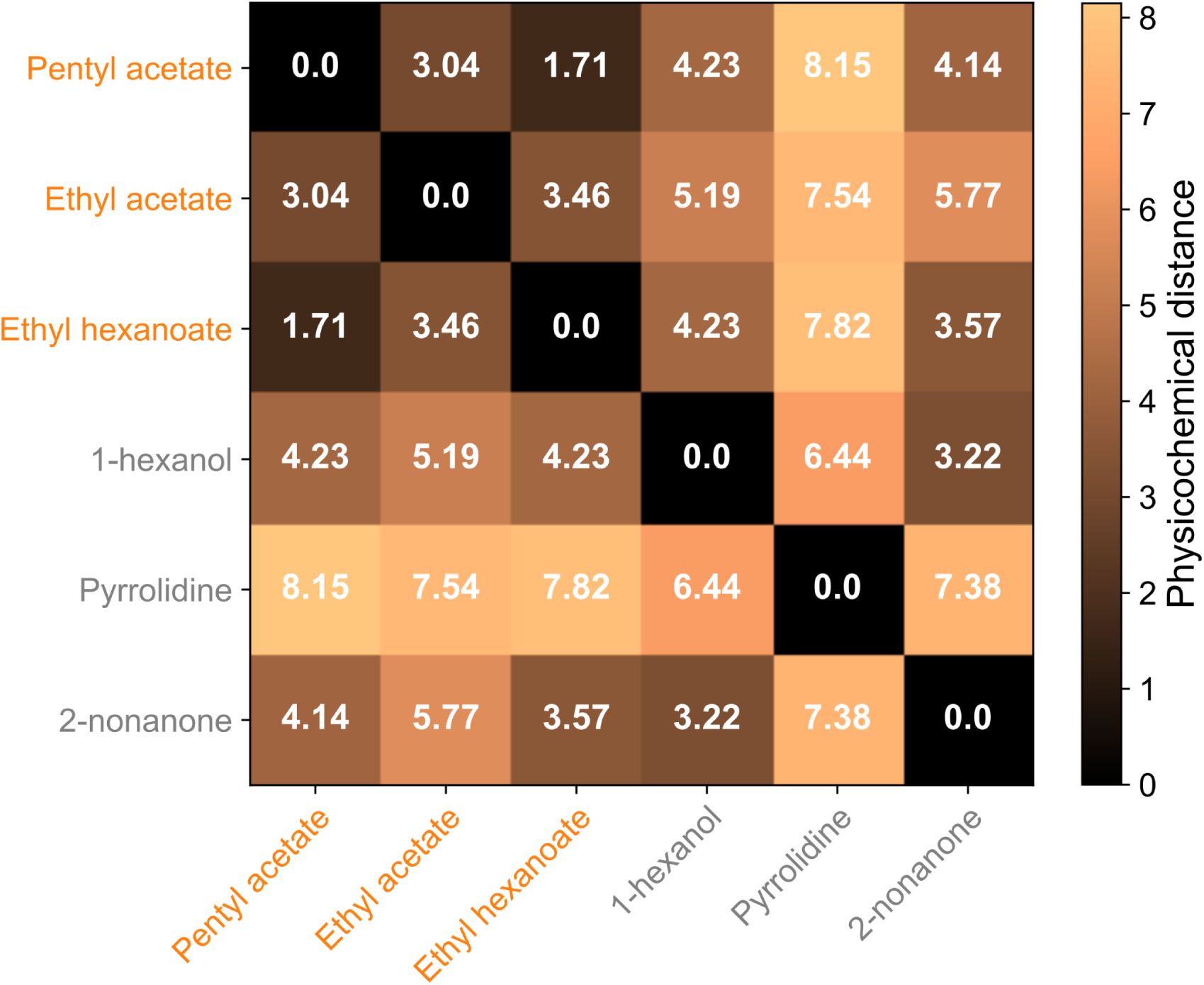
Physicochemical distances between odours. Physicochemical distances between some odours in the experimental odour panel based on Haddad et al. (2008). Food-related odours are shown in orange and danger-related odours are shown in grey.

**Supplementary Figure 3.**
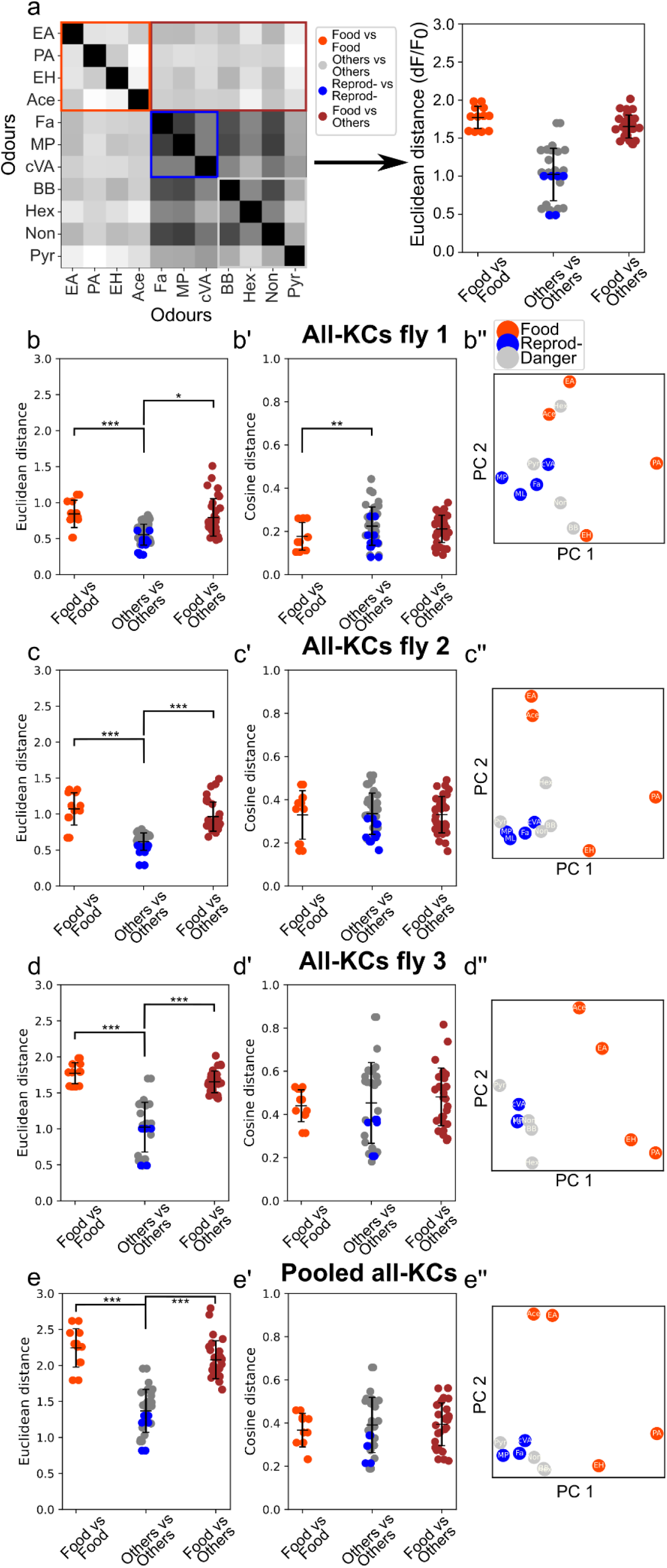
Odour representations in the all-KCs flies. **(a)** Schematic depiction of construction of the distance scatter plot from the corresponding distance matrix. Values in the colored boxes are plotted as points of the same color. The zeros in the diagonal are ignored. Distances between food-related odour representations are shown in orange, between reproduction-related odour representations in blue, between danger-related odours in grey, and between food and other odour representations in brown. * p<0.05, ** p<0.01, *** p<0.001, Kruskal-Wallis H-test. Error bar = Mean ± STD. **(b) - (e)** Euclidean distances between odour representations. **(b’) - (e’)** Cosine distances between odour representations. **(b’’) - (e’’)** Principal component analysis of KC responses to the experimental panel of 12 odours. Odour categories were coloured accordingly: danger (grey), reproduction (blue), and food (orange).

**Supplementary Figure 4.**
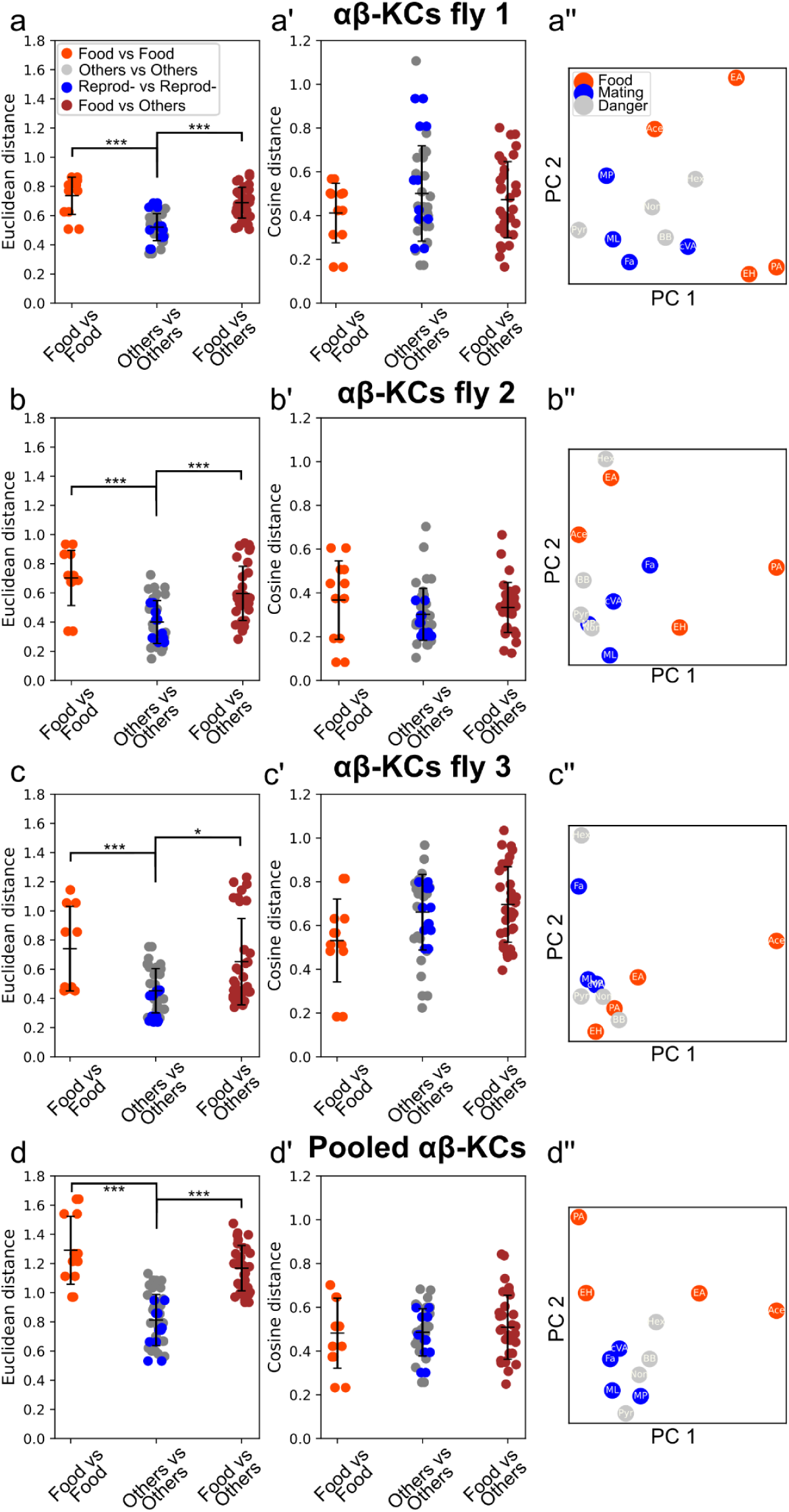
Odour representations in the αβ-KCs flies. **(a) - (d)** Euclidean distances between odour representations. Distances between food-related odour representations are shown in orange, between reproduction-related odour representations in blue, between danger-related odours in grey, and between food and other odour representations in brown. * p<0.05, ** p<0.01, *** p<0.001, Kruskal-Wallis H-test. Error bar = Mean ± STD. **(a’) - (d’)** Cosine distances between odour representations. **(a’’) - (d’’)** Principal component analysis of KC responses to the experimental panel of 12 odours. Odour categories were coloured accordingly: danger (grey), reproduction (blue), and food (orange).

**Supplementary Figure 5.**
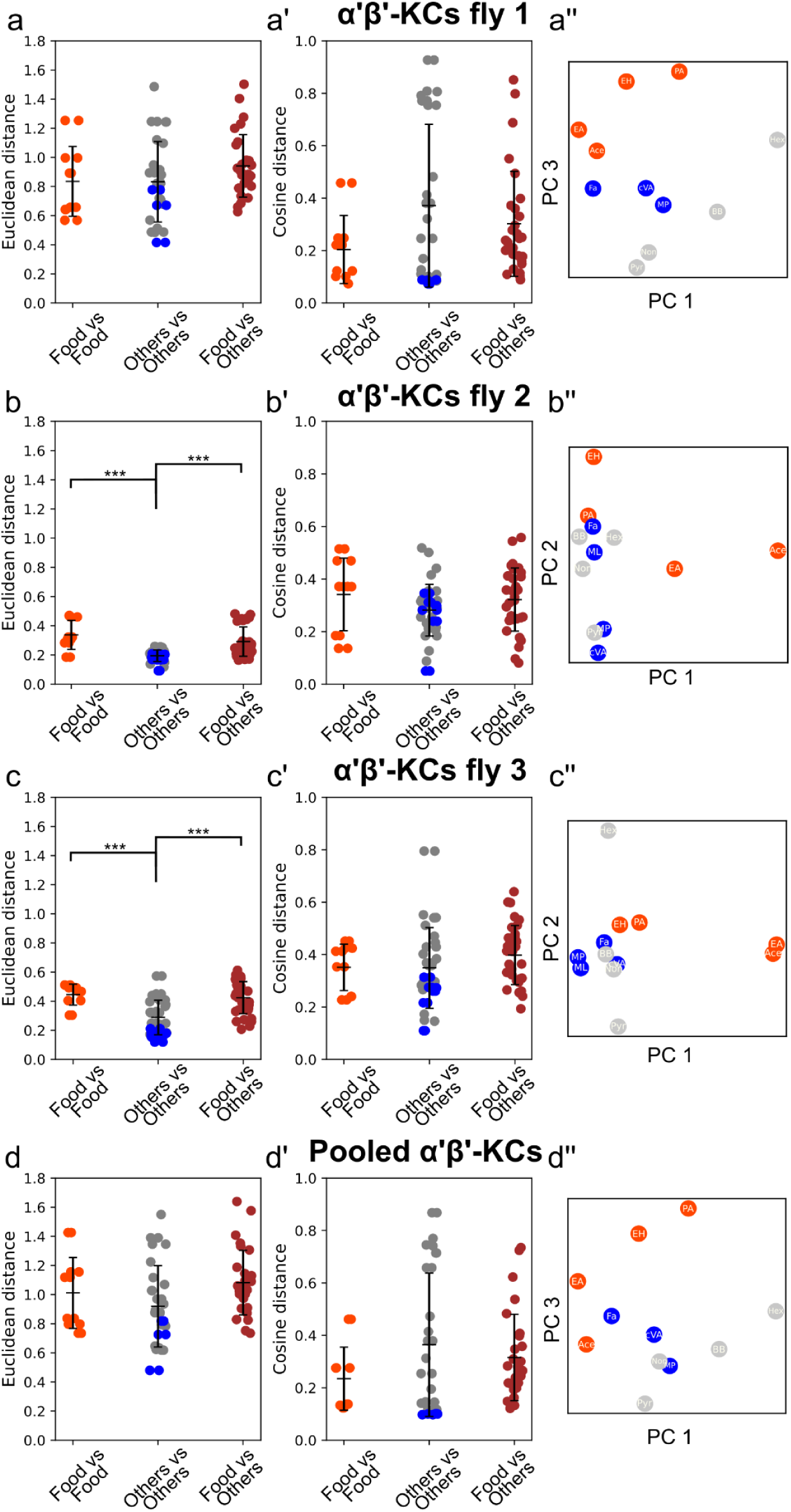
Odour representations in the α’β’-KCs flies. **(a) - (d)** Euclidean distances between odour representations. Distances between food-related odour representations are shown in orange, between reproduction-related odour representations in blue, between danger-related odours in grey, and between food and other odour representations in brown. * p<0.05, ** p<0.01, *** p<0.001, Kruskal-Wallis H-test. Error bar = Mean ± STD. **(a’) - (d’)** Cosine distances between odour representations. **(a’’) - (d’’)** Principal component analysis of KC responses to the experimental panel of 12 odours. Odour categories were coloured accordingly: danger (grey), reproduction (blue), and food (orange).

**Supplementary Figure 6.**
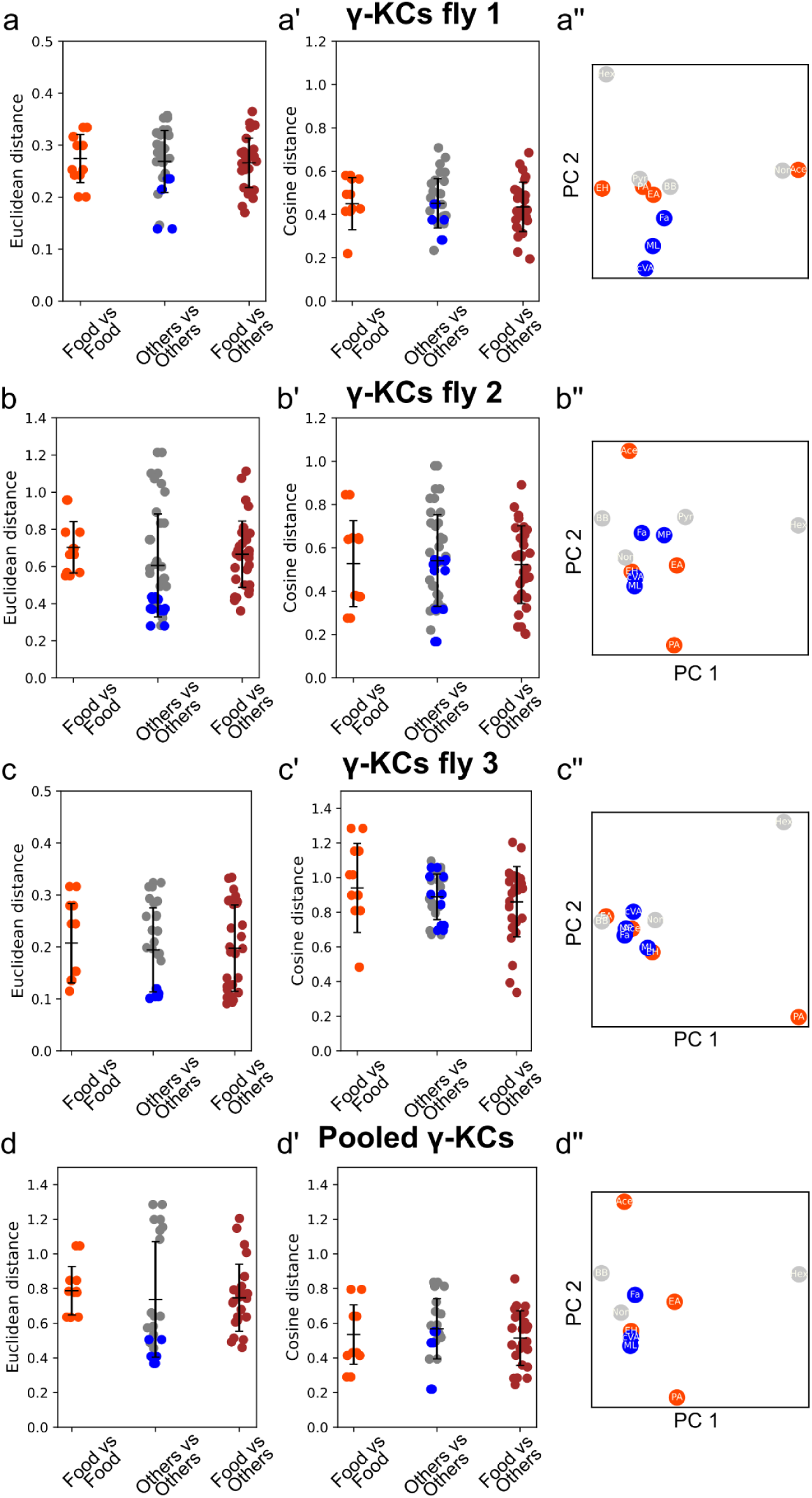
Odour representations in the γ-KCs flies. **(a) - (d)** Euclidean distances between odour representations. Distances between food-related odour representations are shown in orange, between reproduction-related odour representations in blue, between danger-related odours in grey, and between food and other odour representations in brown. * p<0.05, ** p<0.01, *** p<0.001, Kruskal-Wallis H-test. Error bar = Mean ± STD. **(a’) - (d’)** Cosine distances between odour representations. **(a’’) - (d’’)** Principal component analysis of KC responses to the experimental panel of 12 odours. Odour categories were coloured accordingly: danger (grey), reproduction (blue), and food (orange).

## Notes

### Competing Interest Statement

The authors have declared no competing interest.

